# CAdir: Fast Clustering and Visualization of Single-Cell Transcriptomics Data by Direction in CA Space

**DOI:** 10.1101/2025.03.14.643234

**Authors:** Clemens Kohl, Martin Vingron

**Affiliations:** Max Planck Institute for Molecular Genetics, Department of Computational Molecular Biology, Ihnestraße 63-73, 14195 Berlin, Germany

## Abstract

Clustering for single-cell RNA-seq aims at finding similar cells and grouping them into biologically meaningful clusters. Many available clustering algorithms however do not not provide the cluster defining marker genes or are unable to infer the number of clusters in an unsupervised manner as well as lack tools to easily determine the quality of the label assignments. Therefore, clustering quality is commonly evaluated by visually inspecting low-dimensional embeddings as produced by e.g. UMAP or t-SNE. These embeddings can, however, distort the true cluster structure and are known to produce radically different embeddings depending on the chosen hyperparameters. Determining clustering quality therefore still heavily relies on domain knowledge to assess if cells should be clustered together. In order to improve the interpretability of clustering results, we developed CAdir (https://github.com/VingronLab/CAdir), a clustering algorithm that can infer the number of clusters in the data, determine cluster specific genes and provides easy to interpret diagnostic plots. CAdir exploits the geometry induced by correspondence analysis (CA) to cluster cells as well as cluster associated genes based on their direction in CA space. Using the angle between the cluster directions, it is able to automatically infer the number of clusters in the data by merging and splitting clusters. A comprehensive set of diagnostic and explanatory plots provides users with valuable feedback about the clustering decisions and the quality of the final as well as intermediary clusters. CAdir is scalable to even the largest data set and provides similar clustering performance to other state-of-the-art cell clustering algorithms in our benchmarking.

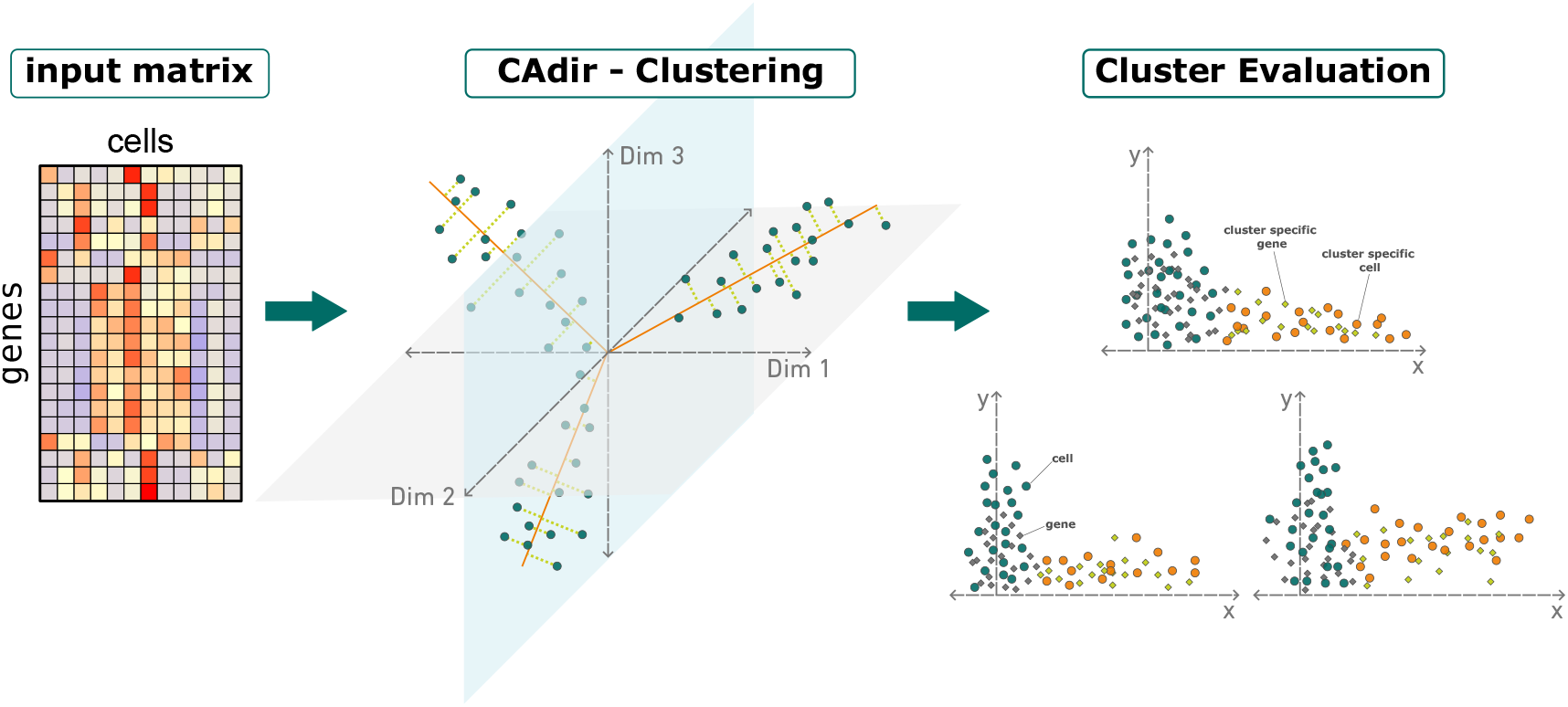

## 1 Introduction

Single-cell RNA sequencing (scRNA-seq) has allowed researchers to study the heterogeneity present in their data and to better understand how different cell types behave. Yet, without prior knowledge and experimental validation, it is often difficult to determine which cell types are present in the data. This is further compounded by the fact that there is no clear definition of a cell type and researchers are often interested in different levels of granularity. Clustering algorithms aim to solve this by unbiasedly grouping cells into biologically meaningful clusters based on their gene expression profiles, thereby helping to discover previously unknown cell types.

Many clustering approaches were developed, including k-means, hierarchical clustering, community detection, and density estimation based approaches [1]. Despite the large number of clustering algorithms available, only a fraction is capable of determining the cluster defining marker genes or estimating the number of cell types in the data [1, 2]. Given the importance of the task, a number of different approaches has been developed to determine the number of clusters, the majority of which use approaches based on intra-/inter-cluster similarity, modularity, stability metrics or eigenvector metrics [2]. Popular approaches for scRNA-seq such as Seurat [3] or Monocle3 [4] use k-Nearest-Neighbour graphs to unfold the high dimensional manifold on which the data lies and then employ graph clustering approaches, e.g. Leiden [5], to find communities in the data without the need to specify the number of clusters *a priori*. While this modularity based approach is able to robustly deal with large data sets, its decision-making process is often opaque to the user and lacks intuitive ways to evaluate the assigned cluster labels.

In order to evaluate the quality of a clustering, researchers therefore often rely on two dimensional embeddings of their data, such as t-distributed stochastic neighbor embedding (t-SNE) [6] or Uniform Manifold Approximation and Projection (UMAP) [7] plots. However, these heuristic methods depend on a large number of hyperparameters and as reported by Chari & Pachter [8], the prediction of a cell’s label based on its k-Nearest-Neighbours is consistently worse using UMAP coordinates than it is with PCA coordinates. Furthermore, UMAP does not take the local density of points into account, resulting in misleading visualizations [9]. For the interpretability of clustering results, the fact that distances in a UMAP or t-SNE cannot be interpreted is particularly problematic, as it often leads to erroneous conclusions drawn from visualizations of e.g. two neighbouring clusters in a UMAP [8].

It is generally recommended not to use UMAP coordinates for clustering due to their stochastic nature and dependency of hyperparameter choices. It therefore stands to reason that they should equally not be used to determine a clustering’s quality, particularly as it can be difficult to understand *why* an embedding created by UMAP, t-SNE or other non-linear methods looks a specific way. If the goal however is to draw a conclusion from the visualization, an interpretable embedding is paramount. Despite their reported shortcomings, UMAP and related methods such as t-SNE have become staple tools used for determining the quality of a clustering due to a lack of easy to use alternatives.

Here we present Correspondence Analysis directional clustering (CAdir), a clustering algorithm that co-clusters cells and genes by their direction in correspondence analysis (CA) [10] space. CA arranges points within a simplex and places points with similar poperties along the same direction. CAdir exploits this property of CA not only to cluster both the cells and genes, but also to estimate the number of groups present in the data. Using the angle between cluster directions as a measure of similarity, CAdir is able to infer when a cluster should be split or merged with another cluster. This dynamic creation and removal of clusters allows CAdir to determine the number of clusters without any prior knowledge about the data.

Unlike other clustering approaches, CAdir offers an intuitive and easy to interpret approach that enables users to evaluate the clustering performance through diagnostic plots and aids in understanding the intermediate steps that lead to the final result. The co-clustered genes further provide the basis for annotating clusters and identifying cluster defining marker genes.

In this paper we introduce the novel clustering approach of CAdir and show that it’s performance is as good as other state-of-the-art cell clustering algorithms, while providing easy to interpret diagnostic outputs, a clustering of both cells and genes as well as determining the number of clusters. Using CAdir’s powerful plotting functionality, we further show how the output of CAdir can be used to better understand and interpret the clustering results.

## 2 Materials and Methods

### 2.1 Correspondence Analysis

Correspondence analysis (CA) is a matrix embedding method similar to PCA. As described by Greenacre [10], the contingency table **P**, which represents the empirical probability that a gene *i* is expressed in cell *j*, is derived by dividing each element of the preprocessed *m* × *n* input matrix *X* with *m* genes and *n* cells by the total sum *n*_++_ of all entries:

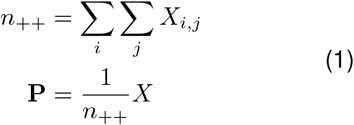

Then, the Pearson residuals **S** are calculated as the deviation of the expected probability *e*_*ij*_ from the observed probability *p*_*ij*_:

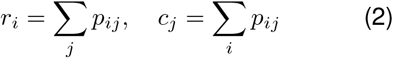

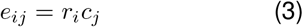

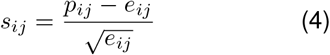

The Pearson residuals are then decomposed using Singular Value Decomposition (SVD):

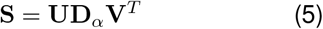

#### 2.1.1 Scaling of the coordinates

In order to obtain interpretable relationships between cells and genes in CA, it is necessary to scale the singular vectors correctly [11]. The standard coordinates are the singular vectors weighted by the row or column masses. With **D**_*r*_ = *diag*(*r*) and **D**_*c*_ = *diag*(*c*) the standard coordinates for genes **Φ** and cells **Γ** are therefore:

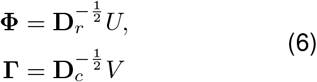

Scaling the standard coordinates furthermore by the singular values, we obtain the principal components **G** of cells and **F** of genes:

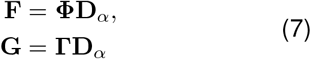

Scaling the cells in standard coordinates **Γ** and genes in principal coordinates **F** provides us with a so called *asymmetric map*, a joint display of cell and gene points with interpretable distances. Using this scaling, the closer a gene is located to a cell the more associated, or specific, it is for a given cell [11].

### 2.2 Clustering by directions

After performing CA, commonly only the *d* first dimensions with the highest inertia are kept in order to remove noise and reduce the size of the data set. A key property of CA that CAdir is built upon, is that CA arranges cells belonging to similar groups along a direction in the dimension reduced space. CAdir capitalizes on this feature by clustering together cells and genes that lie along the same direction (see Fig. 1a). The algorithm consists of three discrete steps that are alternatingly performed during clustering: Dirclust, Split and Merge. After the initial clustering (Dirclust), CAdir attempts to improve the number of clusters in the data based on the angles between clusters during the Split and Merge steps. The cutoff angle required for the split and merge steps is automatically inferred from the data by CAdir, but can alternatively also be set to a user defined

**Figure 1:**
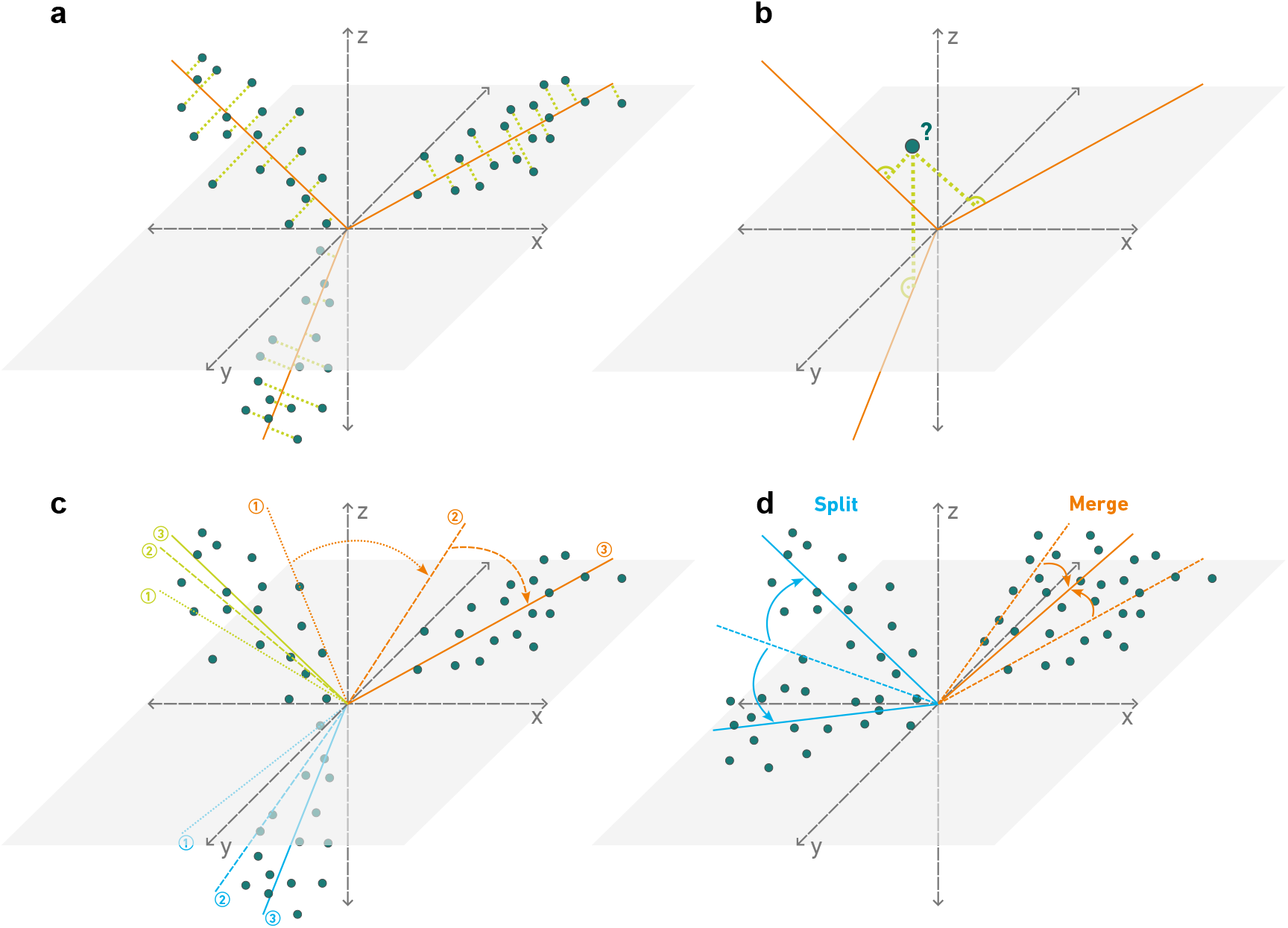
Clustering by direction - Overview. **a**, CAdir clusters cells that lie along a direction in CA space by assigning them to the best fitting line. **b**, Each cell is assigned to the closest cluster direction based on the shortest orthogonal distance of the cell to a direction. The cluster directions are then recomputed based on the changed cluster assignments. **c**, CAdir updates the clusters iteratively: Starting from an initialization (1), cells are assigned to the closest direction and the cluster direction is recomputed (2). This is repeated until the final cluster directions are found (3). **d**, CAdir is further able to independently detect low-quality clusters and thereby infer the number of clusters in the data. Depending on the angle between cluster directions, it either splits a cluster into two new clusters or merges two or more directions into a single cluster. value. CAdir repeats the cluster-split-merge loop several times to ensure that the clustering directions do not change anymore. Lastly, based on the final cluster directions the cluster specific genes are similarly assigned to the closest direction, allowing for cell type discovery and annotation of the clusters.

#### 2.2.1 Detailed description of the algorithm

First during the Dirclust step the *n* × *d* matrix **G** of principal cell coordinates in the *d* dimensional CA space is clustered into *k* initial clusters based on their directions in CA space (see Fig. 1a). The *k* cluster directions thereby form a *L*_*k*×*d*_ matrix, indicating the directions along which the clusters lie in the dimension reduced space. In order to find the shortest orthogonal distance to a direction, the *n* × *k* distance matrix *D* is calculated (Alg. 1, line 8) and each cell is assigned to the closest direction *L*_*k•*_ if the projection of cell *G*_*i•*_ onto direction *L*_*k•*_ is larger or equal to 0 (Alg. 1, line 12 and Fig. 1b). The cluster assignments are stored in the vector *c*.

Given the new cluster members, the cluster directions need to be updated. Total least squares regression is used to find the line that minimizes the orthogonal distance of the cells to the line (Fig. 1a). This can be achieved efficiently by performing SVD on the cells in a cluster. The best fitting line corresponds to the first principal axis 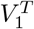 (Alg. 1, line 15 and following line) [12]. Importantly, in order to ensure that points lying on the same line, but opposite directions are not clustered together, the sign of the principal axis is flipped to point into the same direction as the majority of cells in the cluster if the weighted sum *s* in Equation 8 is smaller than 0 [13].

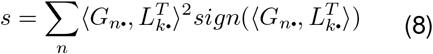

In the pseudocode this is referred to as the `flip_sign` function. The cluster directions are updated for a user specified number of *r* repetitions (usually around 10) (see Fig. 1c).

During the Split step each cluster of cells is split into two sub-clusters using Dirclust with *k* set to two (line 6 in Alg. 2). If the angle of the directions of the sub-clusters is above the pre-defined cutoff angle *θ*, the original cluster is split and the two new sub-clusters are used subsequently, otherwise the original cluster is kept (see Fig. 1d). In order to ensure that a cluster can be split multiple times, the Split function is called recursively after each successful split until all possible splits have been performed (Alg. 2, line 10). Then, the newly split clusters are refined again by calling Dirclust until convergence.

Given, that some of the split-off clusters might actually be part of a larger cluster, CAdir merges clusters during the Merge step. First, the pairwise angles between all directions in *L* are calculated (Alg. 3, line 6). If two or more directions have an angle smaller or equal to the cutoff *θ*, the cells are grouped in a single cluster and a new cluster direction is calculated by determining the first principal axis through SVD (Alg. 3, line 9, Alg. 14 and Fig. 1d). Similar to the Split step, Merge is called recursively after each successful merge to ensure all necessary merges are performed. Finally, the thus newly created clusters are then again refined by calling Dirclust.

##### Algorithm 1

Dirclust

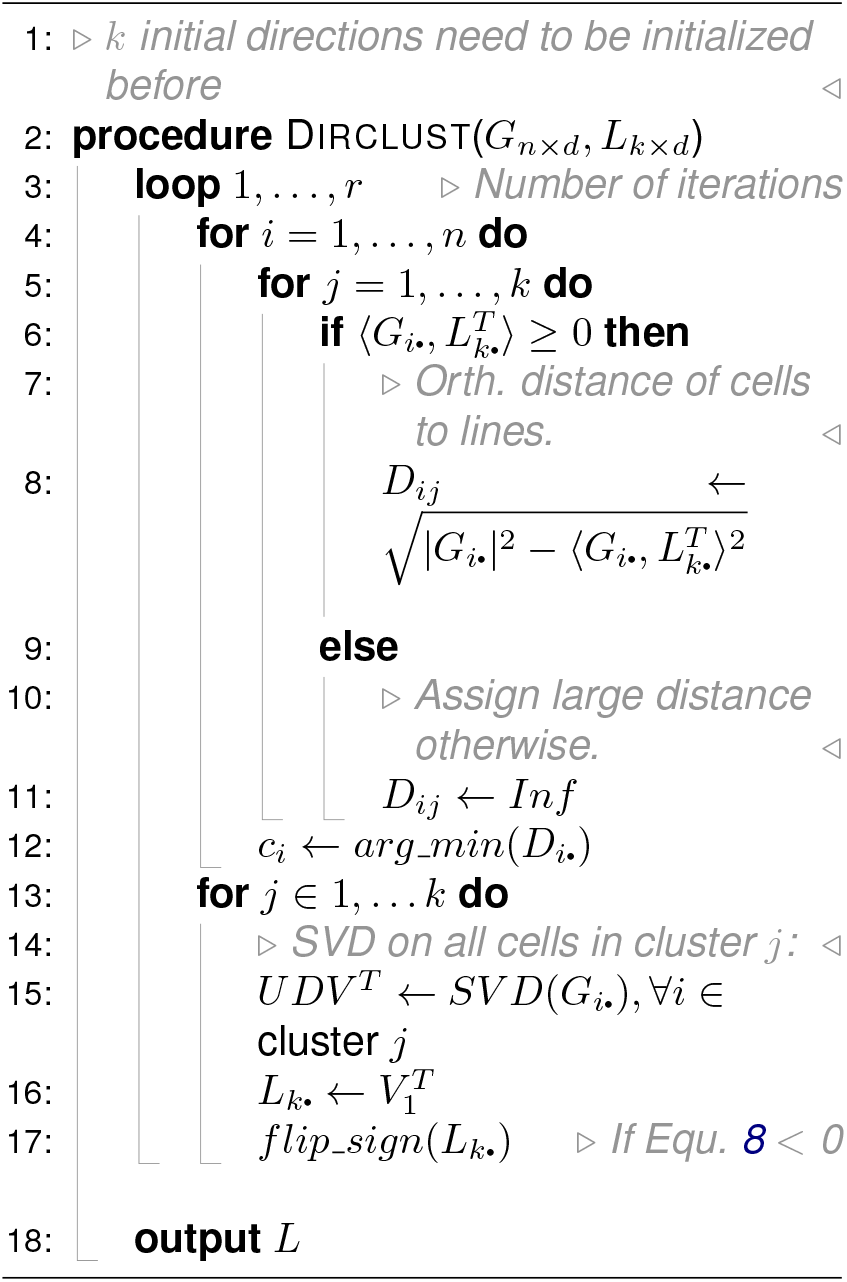

##### Algorithm 2

Split

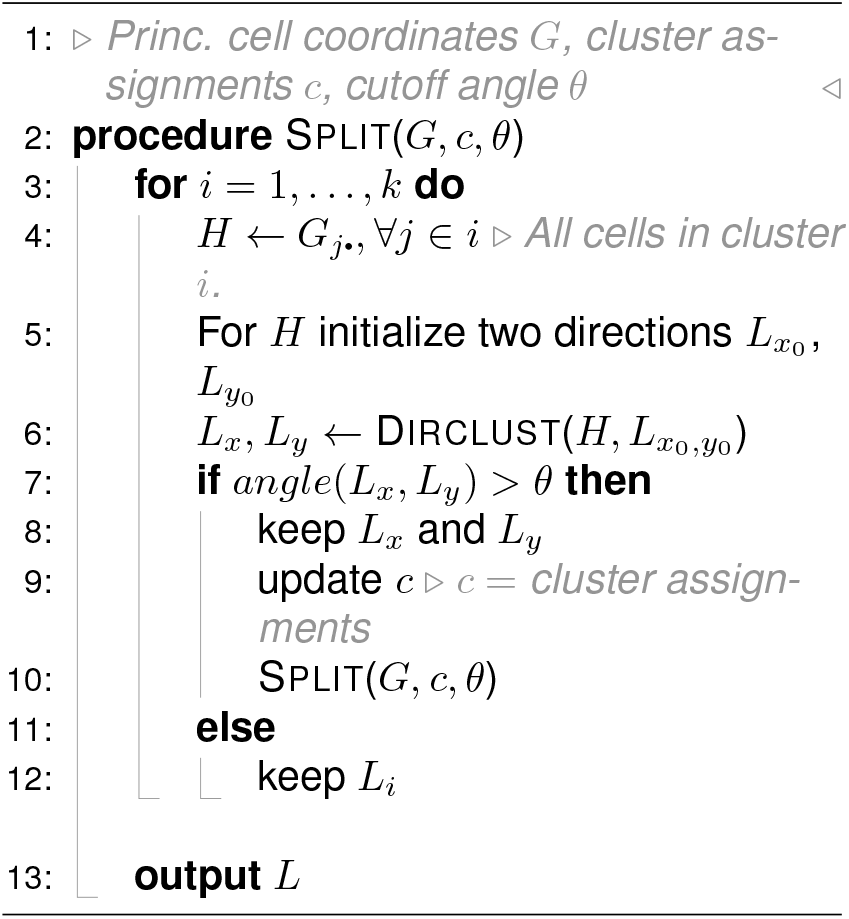

##### Algorithm 3

Merge

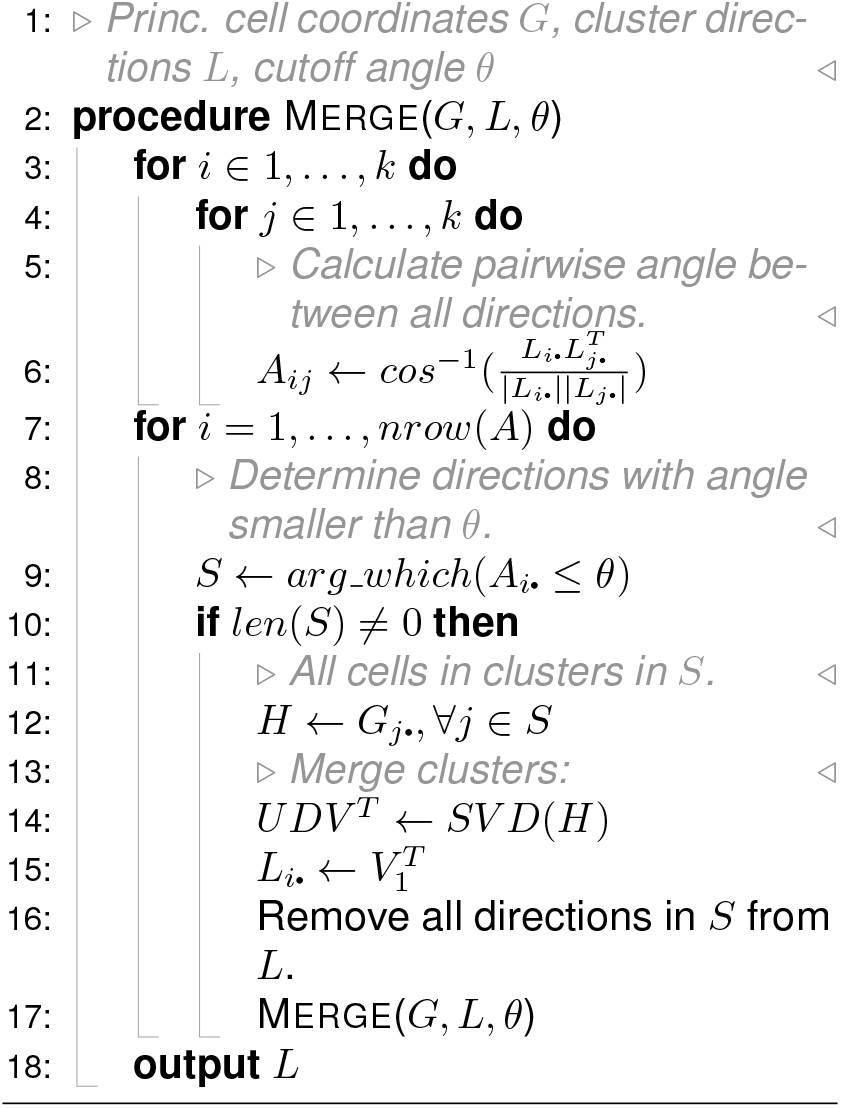

In summary, CAdir can be written as alternating Dirclust, Split and Merge steps as shown in Alg. 4. The initialization (Alg. 4, line 3) of the initial directions *L*_0_ is done by default through k-means_++_ intialization [14], but the directions can alternatively be initialized randomly. After an initial clustering with Dirclust, the directions are split and merged for *r* repetitions. After each Split and Merge step, an additional Dirclust step is added, usually with less repetitions, to refine the cluster directions and to ensure a good fit. The final result is the matrix *L*, whose number of rows (directions) is determined by the algorithm. The cluster assignments are then simply the closest direction for each cell.

### 2.3 Automatic cutoff angle inference

CAdir determines the cutoff angle *θ* for splitting and merging of clusters directly from the data. The angle is inferred by determining the angle above which the majority of cells unrelated to a cluster would fall in an Association Plot [16]. In this publication, this number is determined by many times randomizing the entries in each row (genes) of the input matrix *G* and determining the angle that delineates 99 % of the data. Since this is very time consuming, we have implemented a heuristic which closely approximates the result of this randomization. We select randomly at least 100 directions from the origin and compute the corresponding (random) Associations Plots. Note that due to the random choice of direction we do not expect any meaningful cluster to lie in this direction. The cutoff angle is then set to the angle above which 99 % of the data lie. Typically this angle will be between 60 °and 65 °. In the Association Plot one can sometimes observe that related clusters lie within a much smaller angle.

#### Algorithm 4

CAdir

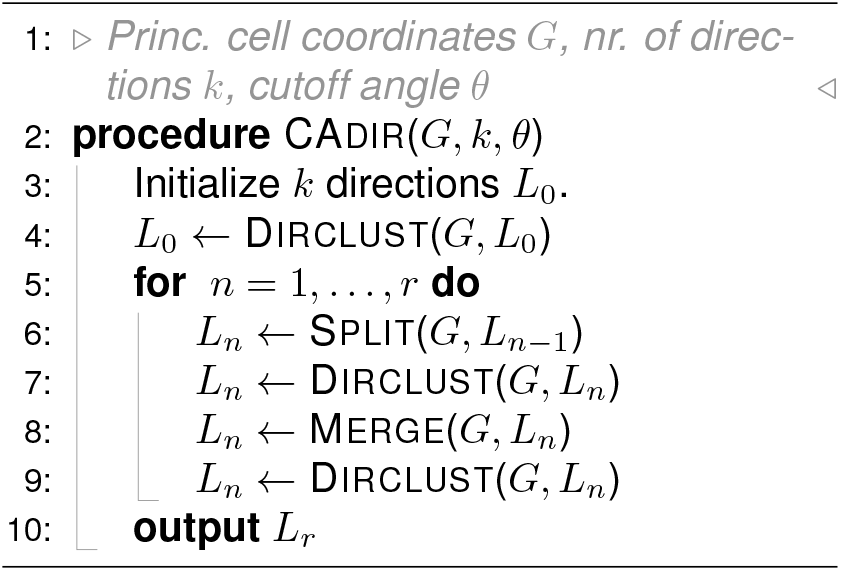

#### 2.3.1 Association Plots

In CA space the cells of a cluster and their specific genes lie along a direction and can be visualized in the form of an Association Plot [15]. The Association Plot is a two-dimensional plot summarizing the position of cells and genes with respect to a direction in space, typically the direction representing the cell cluster. A point represents either a cell or a gene. The x-coordinate of a point in the Association Plot is the distance from the origin to the orthogonal projection of the point onto the direction. The y-coordinate is the orthogonal distance of the point to the direction. For illustration refer to Fig. 1a. There the points in the (high-dimensional) CA space are shown together with their orthogonal projections onto the direction of each of the three clusters. One Association Plot would describe one cluster, with the respective distance to the projection on the x-axis and the orthogonal distances in the y-axis.

The value of Association Plots lies in the interpretability of cluster quality and the relationship between clustered cells and their specifically expressed genes. A well-defined cluster of cells will form a cloud of points extending near the x-axis to the right. Likewise, the specific genes will lie towards the right and close to the x-axis. This is a reflection of the feature of correspondence analysis whereby a cluster of cells with its specific genes extends along a direction away from the origin. Conversely, when a cluster is of low quality, its cells will remain close to the origin and will not be pointed to the right, or they will point upwards rather than follow the x-axis. Correspondingly, there will be few genes to the right and close to the x-axis. Since CAdir clusters the data into directions with associated cells and genes, we utilize the respective Association Plots to visualize clustering results. When CAdir produces *k* clusters, we output *k* Association Plots.

### 2.4 Gene Assignment and Cell Type Annotation

In CA space, genes that are more specific to a cell cluster get placed further out in the direction of the cluster. This feature can be used to assign genes to cell clusters, based on the same criterion of orthogonal distance to the cluster direction. The co-clustered genes are then the input for a gene set overrepresentation analysis using the CellMarker 2.0 [17] gene set to annotate the clusters. For a more detailed description of the gene assignment and the cell type annotation see also the Supplementary Materials.

### 2.5 Datasets and Benchmarking

A detailed description of the used data sets and pre-processing steps, as well as the settings used for benchmarking and evaluation of the results can be found in the Supplementary Materials and in Suppl. Table 1 and 2.

## 3 Results

### 3.1 Clustering cells by direction

Since in CA space clusters of genes and their specific genes get placed along a direction from the origin, our CAdir algorithm looks for distinct, densely populated directions in CA space. Figure 1 shows the basic steps of CAdir as explained in the Methods section. After initializing a pre-defined number of directions, CAdir assigns each cell to the closest direction (see Fig. 1a,b) and re-computes the direction for each cluster through total least squares regression. This is repeated for a set number of iterations until a stable first clustering is reached (Fig. 1c). Then, cluster directions with small angles to each other are merged, whereas sub-clusters with large angles within a cluster are split (Fig. 1d). CAdir determines by itself the number of clusters in the data based on a cutoff angle inferred from the data. The output of CAdir is a set of Association Plots, each visualizing a cluster of cells with its specific genes. This allows for intuitive interpretation of the individual cluster qualities. Based on the cluster specific genes CAdir provides a mapping of clusters to cell types from a given gene set library.

#### 3.1.1 Tabula Muris Limb Muscle data

We clustered the Tabula Muris Limb Muscle (LM) data [20] with 1882 cells using CAdir and used the co-clustered genes to annotate the cells using the automated cell type annotation. As can be seen in Fig. 2a, CAdir finds 7 directions that summarize the data and correspond well to the provided ground truth clustering (SFig. 1c).

**Figure 2:**
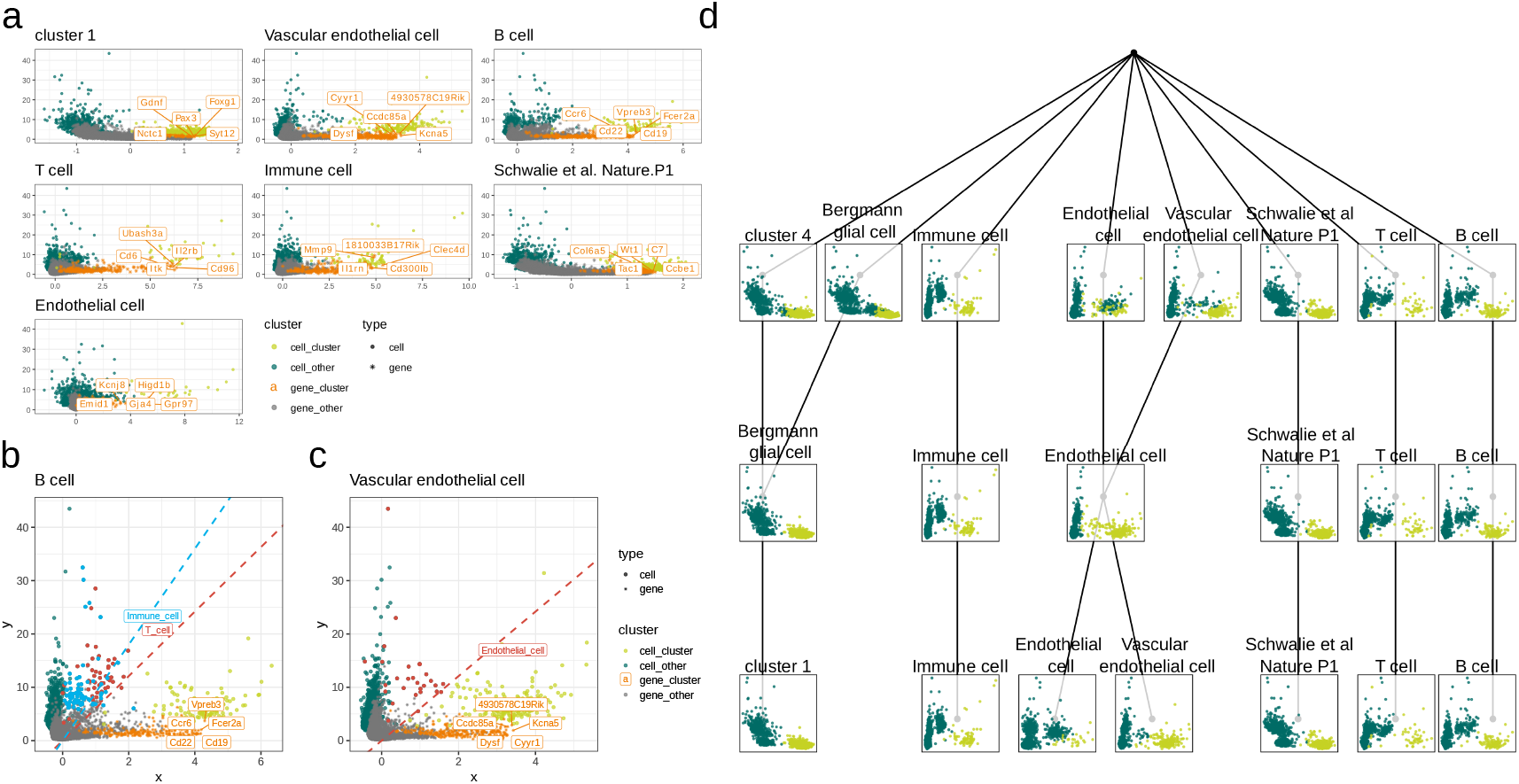
Clustering of Tabula Muris Limb Muscle data. **a**, Association Plots of all clusters determined by CAdir with cluster annotations obtained by gene set over-representation analysis. Clustered cells are colored in lime green and co-clustered genes in orange. Other cells and genes are colored in dark green and grey respectively. For each cluster, the top 5 most highly ranked genes by S_*θ*_-score are labelled. Association plots for **b**, the B cell cluster with the directions of Immune cells (blue) and T cells (red) and the associated cells highlighted and **c**, Association Plot for the Vascular endothelial cell cluster with the direction and the cells of the Endothelial cell cluster highlighted in red. **d**, Graph of the splits and merges performed by CAdir. Each row in the graph corresponds to a single iteration of the algorithm (iterations without splits and merges are removed) and each node to a cluster. At each node an association plot of the cells (principal coordinates) is shown with clustered cells in lime green and all other cells in dark green. The clusters are annotated by the automatic cell type annotation using the co-clustered genes.

CAdir provides visualization tools to comprehensive judge the quality of the obtained clusters: The Association Plots in Fig. 2a are drawn for each cluster, with clustered cells in lime green, co-clustered genes in orange and the remaining cells and genes in dark green and grey respectively. Generally, in a well separated cluster the direction-determining cells have a large x-coordinate in the Association Plot, which corresponds to their projection onto the line. Simultaneously, a low y-coordinate (distance to the line) is desirable, as it indicates that the cluster is not very dispersed and is not, or only mildly, correlated with other directions. For example, the T-cell cluster in Fig. 2a is apparently well defined and, interestingly, so is the unannotated Cluster 1. Although well defined as a cluster, the automatic annotation is undecided here but can be resolved with respect to the annotation of the experiment (Suppl. Fig. 1c).

The angles between cluster directions can also be seen in an Association Plot and this feature can be used to better understand the relationship between clusters. For example, a zoom into the B cell Association Plot shows a low angle between it and the Immune cell and T cell cluster (Fig. 2b and SFig. 1d), as would be expected based on their largely similar expression profiles. Similarly, the projection of the Endothelial cell cluster direction into the Association Plot has a lower angle than other cell types to the Vascular endothelial cells (Fig. 2c and SFig. 1e). The angle between clusters can therefore be used as an indicator of similarity between clusters. The heat map in SFig. 1a shows the angle between cluster directions and SFig. 1b displays this graphically in terms of the directions of a cluster in another cluster’s Association Plot. There one sees, e.g., that there is some relationship between endothelial and vascular endothelial cells, but no similarity between those and the immune cells.

For many applications it can be important to know why a cluster was merged or split. The graph in Fig. 2d visualizes the clustering decisions of the algorithm: Starting from the unclustered data as the tree root, each level represents a split or merge performed by CAdir. The intermediate clustering steps as well as the final clustering were annotated using the method described in section 2.4. To further improve the interpretability, an Association Plot depicting the state of a cluster at a given iteration gives valuable feedback about why it was split, merged, or left as is.

Throughout the iterative process, CAdir combined two clusters into cluster 1 and additionally merged and subsequently re-split the two endothelial cell clusters (Vascular endothelial cell and Endothelial cell respectively). This seemingly redundant step can help to find a better direction for both clusters if the initial clustering included cells from both cell type. By first merging the clusters, CAdir can break out of a local minimum and attempt to find directions that better explain the data.

#### 3.1.2 Determining low-quality clusters

To demonstrate the use of CAdir for quality control and how to identify low-quality clusters, we clustered the PBMC3k data set [21] using CAdir. The clustering results are shown in Fig. 3a. While most clusters show a large number of cluster specific cells and genes, cluster 5 is dominated by only a small number of cells and genes towards the right of the plot (Fig. 3a). The majority of other cells and genes in cluster 5 do not seem to be strongly associated with the cluster and can be found close to the origin. Cluster 5 consists of only a small number of cells (20 cells), and could not be annotated by our automatic cell type annotation.

**Figure 3:**
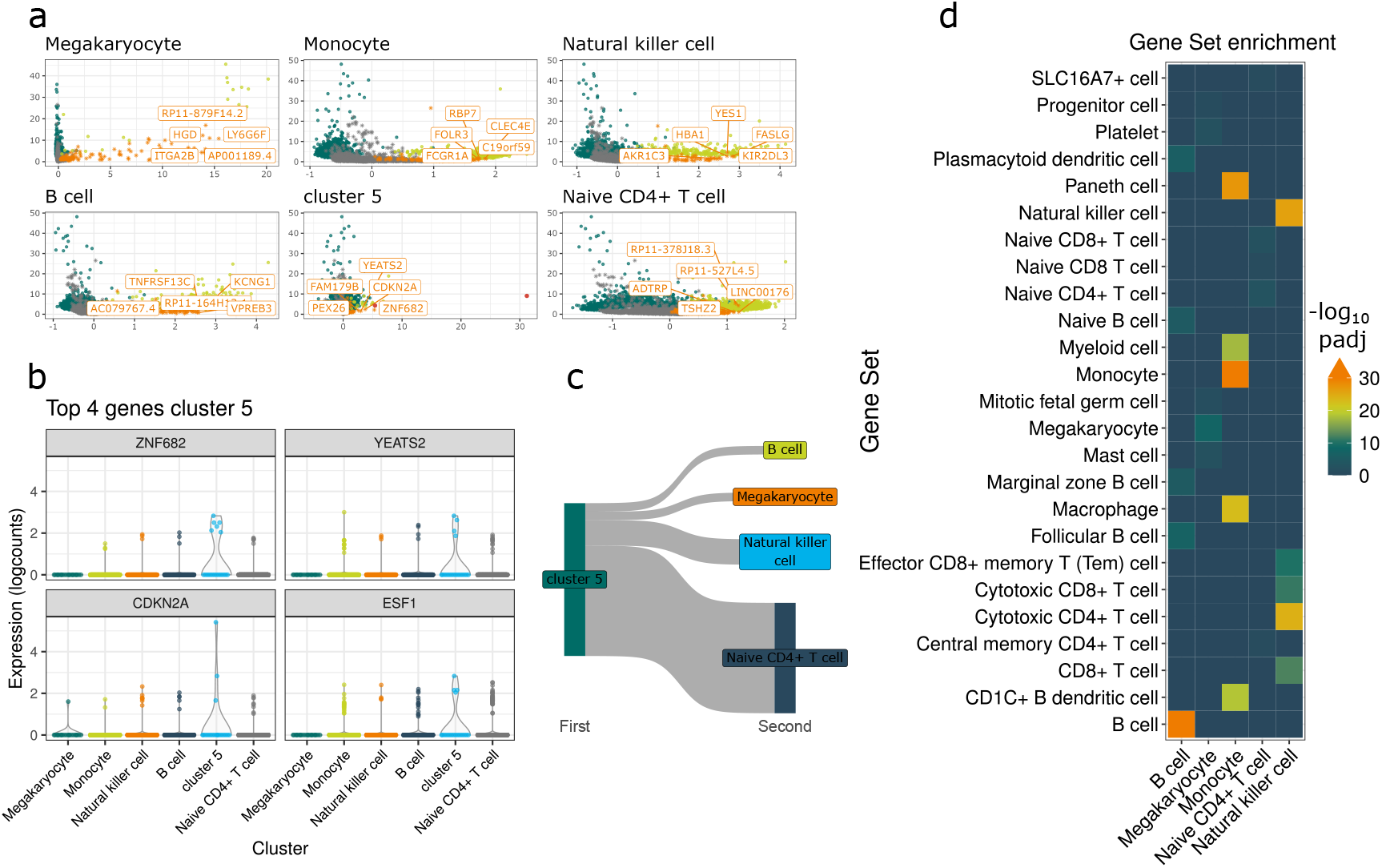
Clustering results for PBMC3k data. **a**, Association Plots for the 6 clusters obtained with CAdir. Clustered cells are colored in lime green, co-clustered genes in orange. Other cells are colored in dark green and the remaining genes in grey. The 5 genes with the highest S_*θ*_-score for each cluster are labelled. The two cells identified as outliers in cluster 5 are marked in red in the respective association plot. **b**, Violin plots of the log-expression of the four genes with the highest S_*θ*_-score for cluster 5. **c**, Sankey plot showing the re-assignment of the remaining cells from cluster 5 after re-clustering to other clusters. **d**, Results of the gene set over-representation analysis used to annotate clusters. The heatmap shows the negative log_10_ adjusted p-values. In order to provide better visual interpretability of the results, the color scale is capped at 30 and gene sets with an adjusted p-value lower than 10^*−*30^ are therefore all colored in orange.

Ranking the genes using S_*θ*_-scores [15], we identified the 4 genes that are most highly associated with cluster 5: ZNF682, YEATS2, CDKN2A and ESF1 (see Fig. 3b). As can be seen in Fig. 3b, most of these genes are expressed only moderately even in cluster 5. ZNF682, which is considerably higher expressed in cluster 5, is predicted to be involved in transcription regulation *ZNF682 Zinc Finger Protein 682 [Homo Sapiens (Human)] - Gene - NCBI* [22], but not known to be indicative of a cell type. The gene CDKN2A, however, codes for 2 proteins, p16^INK4a^ and p14^ARF^, both of which cause cell cycle arrest and generally lead to apoptosis or senescence [23, 24]. CDKN2A is particularly highly expressed in some cells of cluster 5, making it likely that the cluster is mostly based on apoptotic and outlier cells.

Ranking the cells in cluster 5 similarly to the genes, we removed all cells and genes with S_*θ*_-scores above 0. In total this includes 2 cells and 13 genes. The removed cells are also marked in red in the Association Plot for cluster 5 in Fig. 3a. After re-clustering the data with the outliers removed, we obtained the same 5 clusters as shown in Fig. 3a, excluding cluster 5 (see also SFig. 2a). The remaining cells that were originally clustered in cluster 5 are merged into the Naive CD4+ T cell, Natural killer cell, Megakaryocyte and B cell clusters (Fig. 3c), further confirming that cluster 5 was dominated by outliers. The final clustering generally corresponds well with the clusters from the annotation (Sankey plot SFig. 2b), but letting CAdir pick the cutoff angle leads to a slightly more conservative clustering. As can be seen in SFig. 2b, the automatic cell type annotation is able to correctly identify the majority cell type of the clusters based on the co-clustered genes.

One exception is the Megakaryocyte cluster, which is labelled as Platelet cells in the ground truth annotation. This can potentially be attributed to the fact that Megakaryocytes produce Platelet cells, which, as they do not have a nucleus, inherit the mRNA from their parent cell. In cases such as this where two cell types share a large number of expressed genes, the automatic cell type annotation can sometimes mislabel clusters. The results from the gene set over-representation analysis presented in Fig. 3d show that the second most significant gene set (tied with Mast cells) for the Megakaryocyte cluster is in fact Platelet cells. Similarly, Fig. 3d shows that some clusters, such as B cells, have a very clear annotation with one gene set having a much lower adjusted p-value, and other clusters such as Naive Naive CD4+ T cell could have two or more gene sets with very similar p-values. This is also reflected in the Association Plot for the cluster: The plot for the Naive Naive CD4+ T cell cluster shows that the clustered cells are less separated from the other cells compared to e.g. the B cell or Monocyte cluster. Inspecting the cell type annotation results can therefore give additional insights about how well defined a cluster is.

### 3.2 Benchmarking

In order to test the performance of CAdir we benchmarked it against 7 other clustering algorithms. Namely, we tested it against k-means [25], CAbiNet [11], RaceID [26, 27], Seurat [3], Monocle3 [4], SC3 [28] and SIMLR [29]. Because both CAdir and RaceID can either be supplied parameters that determine the number of clusters or be run in an automatic mode in which they infer the number of clusters by themselves, we split it into CAdir/RaceID and CAdir auto/RaceID auto to better differentiate between the two modes.

The tested clustering algorithms differ substantially in the their function: CAdir, CAbiNet, RaceID, Seurat and Monocle3 are able to determine the number of clusters by themselves, whereas k-means, SC3 and SIMLR output a pre-specified number of clusters. In most real world scenarios, the number of clusters *k* is not known *a priori*, and algorithms that automatically determine the number of clusters are therefore often preferred. Additionally, during benchmarking it is not possible to test all possible values of *k*, which can impact the performance of the affected algorithms in the benchmarking.

To minimize bias in our benchmarking process, we evaluated each algorithm across 36 parameter combinations. Additionally, we used the 2000, 4000 and 6000 most highly variable genes in our benchmarking, resulting in a total of 108 runs per algorithm and dataset. From these runs we evaluated the best parameter combination (Fig. 4a,b) in order to further minimize the effect from the choice of parameters.

**Figure 4:**
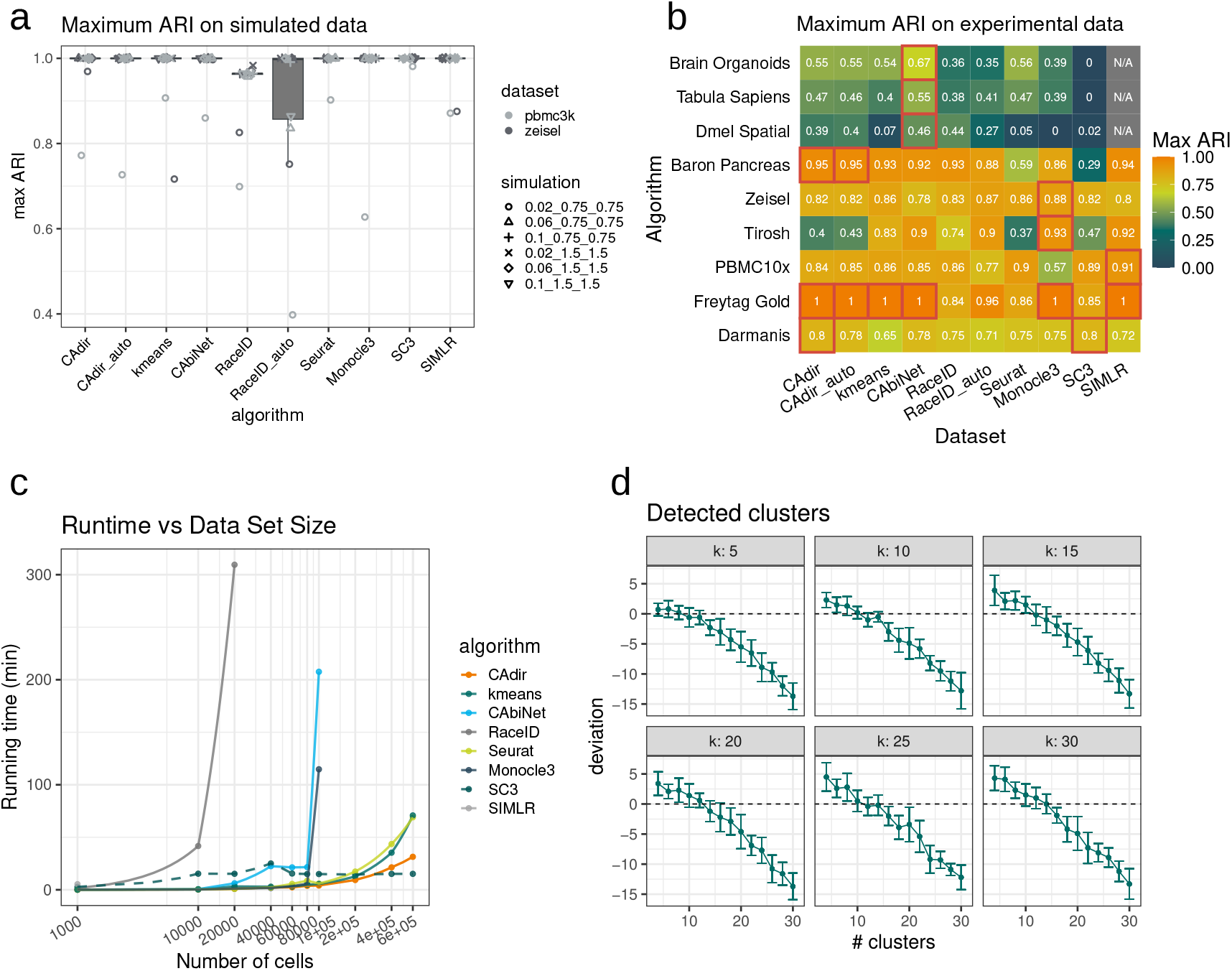
Benchmarking. Highest achieved Adjusted Rand Index (ARI) over all tested parameter combinations for each algorithm on **a**, simulated dataset or **b**, experimental data sets. Only successfully completed runs are considered. The best performing algorithms are highlighted by a red outline in **b. c**, Computational runtime on simulated data of increasing size. SC3 is marked in a dashed line as it is the only benchmarked cluster that does not cluster all cells in a conventional manner but instead only clusters 5000 cells and then attempts to assign the remaining cells to these clusters. **d**, Deviation of the number of retrieved clusters from the number of randomly sampled cell type clusters from the Tabula Muris cell atlas for different numbers of *k* used to initialize CAdir. Error bars indicate the standard deviation over 10 replicate runs.

On simulated data with known ground truth, CAdir performs similarly to other clustering algorithms and even outperforms both RaceID modes (Fig. 4a). For the majority of simulated data sets CAdir obtains an ARI equal to or close to 1, indicating almost perfect correspondence with the ground truth labels. Determining the number of clusters automatically is a challenging task, but while the automated version of RaceID (RaceID auto) performs considerably worse than the version with fixed *k*, CAdir is equally performant when determining the cutoff angle automatically (Fig. 4a). The performance of CAdir, as well as all other tested algorithms, however drops on the two most challenging simulated data sets. Notably, CAdir is extremely robust to the choice of parameters: Using the mean ARI and NMI over all parameter combinations, CAdir on average outperforms all other algorithms except Seurat (SFig. 3a,c).

On experimental data, CAdir is consistently among the best performing algorithms, and on two data sets (FreytagGold and BaronPancreas) even achieves the highest ARI (Fig. 4b). Notably, only Seurat, CAbiNet and CAdir are able to get acceptable clustering results for the two largest data sets, BrainOrganoids and TabulaSapiens. CAdir is able to robustly cluster these large data sets, whereas they often lead to crashes for the algorithms SIMLR, SC3 and RaceID (SFig. 3b). If CAdir tries to infer the cutoff angle from the data (CAdir auto), the maximum achieved ARI is approximately equal to the runs with a predefined angle (Fig. 4b). In some cases, such as the Tirosh data set, it even exceeds the version with a fixed angle.

While CAdir does not outperform all tested algorithms on the simulated and experimental data sets, CAdir provides highly robust clustering results for both small and large data sets. Other algorithms, such as SIMLR, Monocle3 or RaceID, show a significant drop in performance for larger data sets in comparison to the top performing algorithms such as CAbiNet (Fig. 4b). This feature of CAdir is of particular interest due to the continuously increasing size of single-cell RNA-seq data.

Not all clustering algorithms are able to capitalize on their clustering performance when applied to very large data sets such as e.g. tissue atlas data sets, due to long runtimes. CAdir is highly scalable (Fig. 4c) and can cluster a data set with 600 000 cells in approximately 30 min. In our comparison, CAdir is the fastest algorithm tested, even surpassing k-means and Seurat. In particular, SIMLR is unable to cluster any data set with more than 1 000 cells.

Although SC3 has even shorter runtimes, it achieves this by only clustering a small subset of 5000 cells and uses them to train a Support Vector Machine (SVM) to assign the cluster labels to the remaining cells [28]. Due to the small number of cells sampled, this approach however does not adequately represent the diversity present in large scRNA-seq data sets. This is also evident from the poor clustering results for the moderately large BrainOrganoids and TabulaSapiens data sets reported in Fig. 4b.

We furthermore tested the ability of CAdir to correctly identify the number of clusters in the data. Similarly to Yu *et al*. [2], we sampled increasing amounts of cell type clusters from the Tabula Muris cell atlas [30] and tested if CAdir is able to identify the correct number of clusters when estimating the cutoff angle by itself. Up to approx. 20 clusters CAdir provides cluster estimations very close to the ground truth (Fig. 4d). While CAdir is able to recover the correct amount of clusters at the beginning, the deviation from the sampled number of clusters grows with an increasing number of cell types.

Notably, the Tabula Muris data consists of cells from 18 different tissues, many of which contain similar cell types. This leads CAdir to primarily cluster by tissue instead of by cell type, explaining why it performs best around 15-20 sampled clusters.

We additionally explored the effect of the initial *k* on the final results. Testing 6 different choices of *k* to initialize CAdir, the differences between the runs are marginal (Fig. 4d). This further confirms the ability of CAdir to determine the cluster number largely independent of the initialization. Similarly, the choice of of the cutoff quantile used to automatically determine the angle for splits and merges seems to only have a minor effect (SFig. 3d). Generally, a more restrictive quantile cutoff, which results in a lower cutoff angle, appears to slightly improve the recovery of the correct number of clusters.

## 4 Discussion

CAdir is a versatile clustering algorithm that aims to provide actionable information on clustering decisions and the quality of the results. Unlike many other popular scRNA-seq clustering algorithms, CAdir can be run in a fully automatic mode in which CAdir infers the cutoff angle *θ* by itself to determine the number of clusters instead of relying on user defined parameter choices. The angle between the cluster directions is useful beyond just determining whether a cluster should be split or merged: Unlike distances between clusters, the angles between clusters provide an interpretable indicator of similarity that helps to identify similar cell types. Combined with its competitive clustering performance and its fast runtime, CAdir is ideally suited for the analysis of single-cell RNA-seq data. In particular, it is able to cluster very large data sets in a fraction of the time required for other popular tools such as Seurat, SC3 or SIMLR.

The concept of clustering by direction intuitively lends itself to visualizing the clustering results in Association Plots, which provide a two dimensional visualization of the cluster based on its direction in CA space. This method allows for a more direct evaluation of a cluster’s quality due to its interpretable axis compared to e.g. UMAP, which might distort the data in unforeseen ways. Beyond just providing a straightforward and intuitive way to judge a clusters quality, CAdir provides a visualization of the clustering decisions made by the algorithm in the form of a graph tree. The graph view of the splits and merges further facilitates the process of understanding if a cluster appropriately represents a distinct community of cells.

Unlike other algorithms that only provide a clustering of the cells, CAdir also assigns genes that are highly associated with a group to the same cluster. This not only allows researchers to understand which genes define a specific cluster, but enables fast cell type annotation without the need for differential gene expression analysis or other methods.

As a purely linear method, CAdir can struggle to find well fitting lines in the presence of strong non-linearities in the cluster structure. This could for example be the case when analysing developmental data with many differentiating cells. In our benchmarking we have two such data sets, Brain Organoids and the Drosophila spatial data (Dmel Spatial) but in our analysis, CAdir performs en par or even outperforms other clustering algorithms such as Seurat or CAbiNet. Future iterations of the algorithm could be improved by allowing for non-linear cluster shapes through e.g. kernel methods.

In summary, CAdir is a fast and capable clustering algorithm that provides interpretable output and can automatically infer the number of clusters from the data. The R package CAdir can be installed from GitHub: https://github.com/VingronLab/CAdir.

## 5 Code availability

The latest Version of CAdir can be downloaded and installed from GitHub (https://github.com/VingronLab/CAdir). The Version of the package used to produce the results presented in this manuscript is available on Zenodo (10.5281/zenodo.14900875). The code to reproduce the presented results can be found here on GitHub (https://github.com/VingronLab/CAdir_results) and is also deposited on Zenodo (10.5281/zenodo.14894256).

## 6 Data availability

The discussed simulated and experimental data sets can be downloaded from Zenodo: 10.528 1/zenodo.14894256

## 8 Author contributions statement

M.V. and C.K. conceived and conceptualized the algorithm, C.K. and M.V. wrote and reviewed the manuscript, C.K. implemented the method and conducted and analyzed the experiments.

## 9 Acknowledgments

The authors wish to thank the IT group of the Max Planck Institute of Molecular Genetics for their support and thank the anonymous reviewers for their valuable feedback.

