## Supplementary Materials for "CAdir: Fast Clustering and Visualization of Single-Cell Transcriptomics Data by Direction in CA Space"

14[th] March, 2025

#### Supplementary methods and materials

##### Gene Assignment

In an assymetric map in CA, cluster specific genes are located along the cluster direction, similarly to cells belonging to this cluster and similarly to the cells they can therefore be assigned to the cluster based on the orthogonal distance to the cluster direction. The most specific genes lie far away from the origin while around the origin all the unspecific genes are located. Therefore, we filter out those genes that are within a sphere of radius $r$ around the origin. The radius $r$ is chosen to filter out $80\%$ of genes. Only the remaining genes get assigned to the cluster direction with the shortest distance of the gene's standard coordinates to its projection on the line. We found that the genes are best filtered based on their principal coordinates, but the actual cluster assignment is done in standard coordinates in order to ensure interpretable distances. Furthermore, genes get ranked for specificity for their cluster according to the specificity score $S$ as introduced by Gralinska *et al.*:

$$S_\theta = x - \frac{y}{\tan\theta}, \quad (1)$$

where $x$ and $y$ are the coordiantes of a gene in the Association Plot.

##### Cell Type Annotation

The CAdir package allows for automatic cell type annotation using the co-clustered genes. Using the CellMarker 2.0 [**huCellMarkerUpdatedDatabase2023**] gene set, each cluster is annotated as follows: The gene sets are filtered to only contain sets with a minimum of 10 and a maximum of 500 genes and only genes that are present in the single-cell data set and could therefore be assigned to a cluster are kept. A hypergeometric test is then performed for all co-clustered genes in a cluster and all gene sets. Obtained p-values are adjusted to correct for multiple testing and the adjusted p-values are used to construct a cost matrix containing the clusters and all enriched gene sets. If a cluster's genes were not found to be enriched in a specific cluster, its cost gets set to 1. Using the R package RcppHungarian (`https://doi.org/10.32614/CRAN.package.RcppHungarian`), we applied the Hungarian method [**kuhnHungarianMethodAssignment1955**] to solve the assignment problem, such that each cluster is getting uniquely assigned a cell type, even if clusters have similar enriched gene sets. If, however, the assigned gene set has an adjusted hypergeometric p-value smaller than 0.05 the cluster is left unannotated.

#### Association Plots

Association Plots are constructed as described by Gralinska *et al.*. The Association Plot x-coordinate of a point $a$ is calculated as the scalar projection of its vector $\vec{b}$ in CA space onto the cluster direction $\vec{d}$, whearas the y-coordinate is the distance of the point to the direction:

$$\begin{aligned} a_x &= \frac{\vec{b}\vec{d}}{\|\vec{d}\|}, \\ a_y &= \sqrt{\|\vec{b}\|^2 - a_x^2} \end{aligned} \tag{2}$$

#### Data Processing

Simulated and experimental data sets used for the benchmarking were pre-processed as follows: Using the functions `perCellQCMetrics` and `perCellQCFilters` from the R package scuttle [2], we removed cells that are identified as outliers based on the number of detected genes, the sum of counts per cell and the percent of mitochondrial reads. Additionally, we removed genes that are expressed in less than 1 % of cells. Afterwards, cells were normalized and counts were transformed to log-counts using the functions `quickCluster`, `computeSumFactor` and `logNormCounts` from the packages scran [3] and scuttle [2].

The brain organoids data was pre-processed similar to the other data sets, but as was also done in the original publication [4], only cells with less than 40 % mitochondrial counts were kept.

#### Simulated Data

Simulated data was generated using the Bioconductor package Splatter [5], which can generate new single-cell RNA-seq data with arbitrary number of clusters based on parameters estimated from experimental data. We used two different data sets, the Zeisel Brain Data [6] (Zeisel) obtained through the Bioconductor package scRNA-seq [7] and PBMC3k data from 10x Genomics obtained through the Bioconductor R package TENxPBMCData, [8] to estimate parameters concerning the mean gene expression, outliers, library size, the biological coefficient of variation and dropouts.

We used each real data set (Zeisel and PBMC3k) as the basis for 6 different simulated data sets for which we varied the probability for a gene to be differentially expressed as well as the shape and mean parameter of the log-normal distribution that defines the magnitude of the differential expression, resulting in a total of 12 data sets (see Suppl. Table 1). Each simulated data set was generated with 1,000 cells, 10,000 genes and 6 six clusters. The generated clusters contained 30 %, 25 %, 20 %, 10 %, 10 % and 5 % of the total number of cells respectively.

#### Experimental data

Experimental data sets were downloaded directly from the relevant publications or alternatively obtained through R data repositories such as the scRNAseq and TENxPBMCData Bioconductor packages. In our benchmarking we used the following data sets: Freytag Gold [9], Tabula Muris [10], Tirosh [11], PBMC10x [12], Dmel Spatial [13], Tabula Sapiens [14] and Brain Organoids [4] data sets were downloaded from the resources specified in the respective publication. Using the Bioconductor package scRNAseq [7] we downloaded the following data sets: Zeisel Brain [6], Darmanis [15] and Baron Pancreas [16]. The Tabula Muris LM (Tabula Muris subsetted to Limb Muscle tissue) [10] data was downloaded using the TabulaMurisData R Bioconductor package [17] and the PBMC3K data was downloaded using the TENxPBMCData Bioconductor package [8]. See also Suppl. Table 2. All data sets were pre-processed as described in the Suppl. Materials section Data Processing.

#### Discussed data

The Tabula Muris Limb Muscle and PBMC3k data sets discussed in the results were processed as following: We downloaded the Smartseq2 Tabula Muris data with the Bioconductor package scRNAseq [17] and subsetted it to the Limb Muscle tissue. It was processed as described above and then subset to the 6000 most highly variable genes using the R package scran, resulting in a data set with 1882 cells and 6000 genes. We performed CA using the package APL and kept the first 30 dimensions. Subsequently, the data was clustered using CAdir with $k = 8$ and the angle cutoff set to 55 degrees. Cells were annotated by gene set overrepresentation analysis using the CellMarker 2.0 gene set.

The PBMC3k data was downloaded from [8] (`https://doi.org/10.18129/B9.bioc.TENxPBMCData`) and preprocessed according to the Seurat "Guided Clustering Tutorial" [18] (`https://satijalab.org/seurat/articles/pbmc3k_tutorial`, accessed on 2024-08-23). We retained the 20 % most highly variable genes, performed CA and kept the first 30 dimensions. Clustering with CAdir was performed with $k = 12$ and the cutoff angle was inferred automatically by CAdir. The co-clustered genes were used to annotate the data using the included gene set overrepresentation analysis method.

#### Benchmarking

We benchmarked CAdir against 7 other clustering algorithms, namely Seurat [19], CAbiNet [20], Monocle3 [21], SIMLR [22], SC3 [23], RaceID [24, 25] and k-means clustering.

In order to test the algorithms under different conditions, we benchmarked them on the simulated and real data sets using the 2000, 4000, and 6000 most highly variable genes as determined by the function `modelGeneVar` and `getTopHVGs` from the package scran [3]. To prevent any bias in the choice of parameters, we additionally ran every algorithm with 36 different parameter combinations for a total of 108 runs per algorithm and data set. As the number of parameters differ between algorithms, this can result in many variations of a single parameter or only a few different choices for more parameters. The data was pre-processed in the same way for all algorithms prior to clustering. Algorithms that failed during the first run were rerun with more computational resources. If for example an algorithm required more memory than was provided, it was rerun with 500 Gb of memory, but if it exceeded the run time limits, the algorithm was restarted with a run time limit of 24 hours. For any other failures the algorithms were rerun with double the amount resources as for the original run, with a maximum of 500 Gb of memory and 24 hours of runtime. Algorithms that did not finish the clustering were rerun up to two times.

#### Evaluation

We used the Adjusted Rand Index (ARI) [26] and the normalized mutual information (NMI) [27] as implemented in the R package aricode [28] to compare the cell clusterings to the reference annotations. The Adjusted Rand Index measures the similarity between two clusterings as done by the Rand Index, but is adjusted to correct for coincidental clusterings. The ARI ranges from -1 to 1, where 0 indicates a random clustering, -1 a clustering worse than expected by chance and 1 that the two clusterings match perfectly. As the name already implies, NMI is a normalized measure for mutual information, and is bounded between 0 and 1. As it measures the dependence between the two groupings, higher values for the NMI indicate a higher association between the two clusterings.

Only successful runs were used for the evaluation, meaning that the summary statics such as mean or the maximum are taken over differing amount of runs, depending on how many effectively finished. We furthermore excluded runs that did not return interpretable results. Some clustering algorithms for example return "NA" or 0 clusters when they do not converge. These results were therefore excluded.

##### Runtime benchmarking

Similarly to the benchmarking performed for the cell clustering, we used Splatter to simulate data sets of increasing size. We simulated data sets with 2 000 genes and 1 000, 10 000, 20 000, 40 000, 60 000, 80 000, 100 000, 200 000, 400 000 or 600 000 cells. For all simulated data sets we simulated 6 clusters and set the parameters `de.facLoc` and `de.facScale` to 1.5. The probability for a gene to be differentially expressed was set to 0.1. The simulated data sets were normalized using the functions `quickCluster` and `logNormCounts`. All tested algorithms were run using a single thread and the runtime including potential dimensionality reduction steps was recorded.

##### Cluster number recovery

To benchmark how well CAdir recovers the correct number of clusters, we used a scheme similar to the one described by Yu *et al.*: We subset the pre-processed Tabula Muris cell atlas data to cell types with at least 300 cells, resulting in a data set 15 771 genes × 39 207 cells with 38 cell types from 18 tissues. We then subset the data set to $n$ randomly sampled cell types, where $n$ is between 4 to 30 cell types in steps of 2, and again sub-sampled each cluster to 200 cells in order to equalize the size of the clusters. We retained the top 4 000 most highly variable genes, kept the first $n + 20$ dimensions after performing CA and let CAdir automatically infer the cutoff angle below which 99 %, 99.9 % or 99.99 % of cells fall when considering random directions in CA space. To account for the variability introduced by randomly sampling clusters, we repeated each run 10 times.

#### Supplementary Results

### Supplementary Tables

| DE prob. | DE factor mean | DE factor var. | Name |
|---|---|---|---|
| 0.02 | 0.75 | 0.75 | (pbmc3k / zeisel)_0.02_0.75_0.75 |
| 0.06 | 0.75 | 0.75 | (pbmc3k / zeisel)_0.06_0.75_0.75 |
| 0.1 | 0.75 | 0.75 | (pbmc3k / zeisel)_0.1_0.75_0.75 |
| 0.02 | 1.5 | 1.5 | (pbmc3k / zeisel)_0.02_1.5_1.5 |
| 0.06 | 1.5 | 1.5 | (pbmc3k / zeisel)_0.06_1.5_1.5 |
| 0.1 | 1.5 | 1.5 | (pbmc3k / zeisel)_0.1_1.5_1.5 |

**Suppl. Table 1: Parameter combinations for simulated data used for benchmarking.** The 12 simulated data sets used in benchmarking the cell clustering are either based on the PMBC3k or Zeisel data set. For each reference data set we estimated the base parameters and then varied the mean of the log-normal distribution (**DE factor mean**), the variance of the log-normal distribution (**DE factor var.**) and the probability for a gene to be differentially expressed (**DE prob.**).

### Supplementary Figures

| Data Set | cells × genes | Description |
|---|---|---|
| Darmanis | 461 × 17 533 | Human adult cortical tissue |
| Freytag Gold | 914 × 19 973 | Human lung adenocarcinoma cell lines |
| Tabula Muris LM | 1882 × 13 204 | Mouse limb muscle tissue |
| Zeisel Brain | 2874 × 14 508 | Mouse cortex and hippocampus |
| PBMC3k | 2700 × 32 738 | Human PBMCs |
| Tirosh | 2880 × 16 347 | Human melanoma |
| PBMC10x | 3176 × 11 881 | Human PBMCs |
| Baron Pancreas | 8569 × 12 238 | Human Pancreas |
| Dmel Spatial | 14 808 × 7 178 | *D. melanogaster* embryo |
| Tabula Sapiens | 32 393 × 15 674 | Human endothelial cells |
| Brain Organoids | 35 291 × 10 640 | Human cerebral organoids |
| Tabula Muris | 15 771 × 44 104 | Mouse tissue |

**Suppl. Table 2: Overview over the used data sets.** The name listed under **Data Set** is used as the shorthand name for the data set. Cell and gene counts refer to the pre-processed data before feature selection.

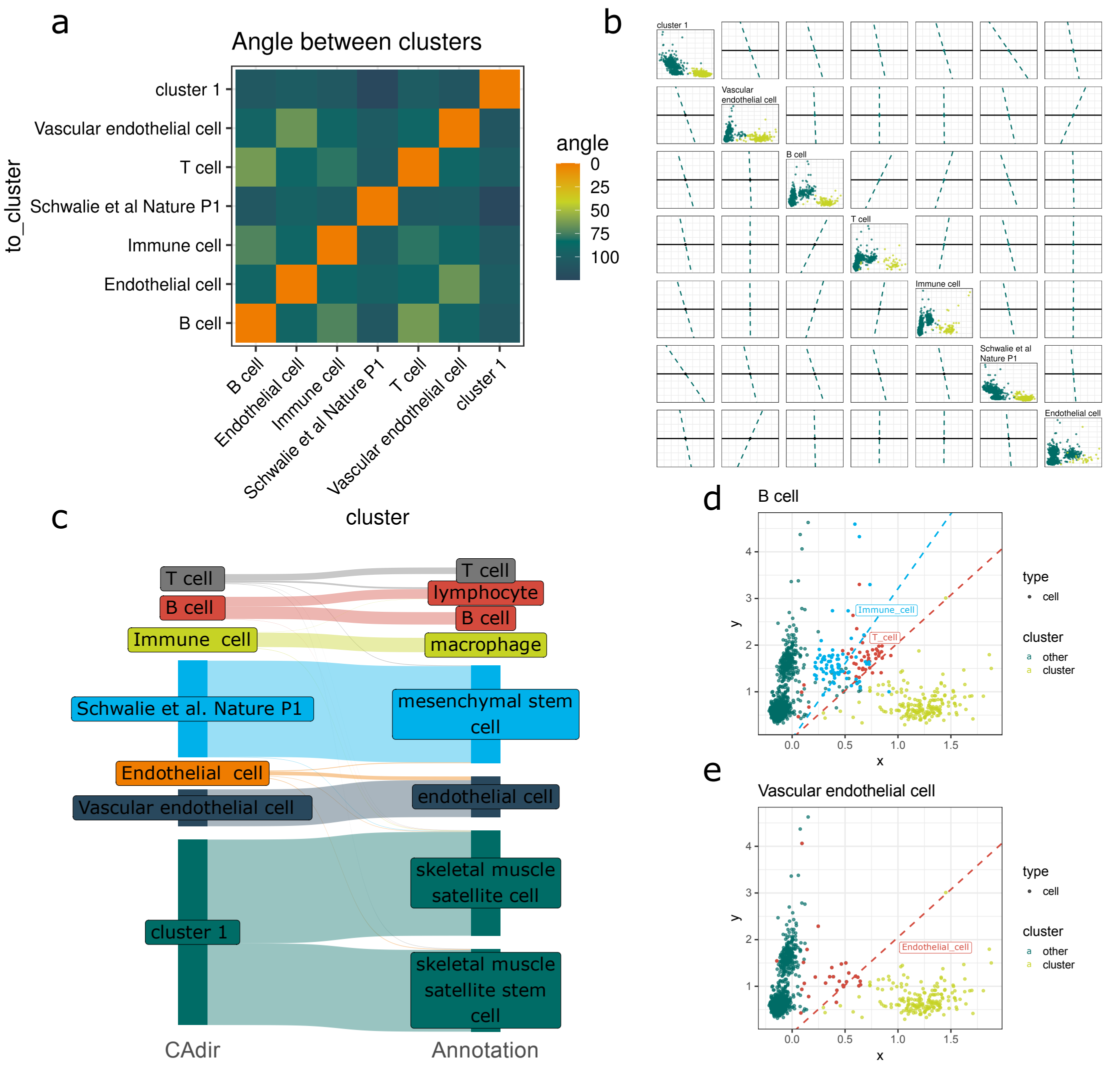


**Suppl. Fig. 1: Interpretation of the Tabula Muris Limb Muscle data clustering.** **a**, Matrix of the pairwise angles between the cluster directions. Lower angles indicate higher similarity between clusters. **b**, Each row shows an Association Plot of the respective cluster in the diagonal and lines that show the direction of the remaining other clusters projected into the Association Plot. **c**, Sankey plot of the annotated clustering obtained with CAdir (left) and the ground truth annotation (right). Association Plot with cells in principal coordinates for **d**, B cells and **e**, Vascular endothelial cells. Closely associated clusters are colored in red and blue.

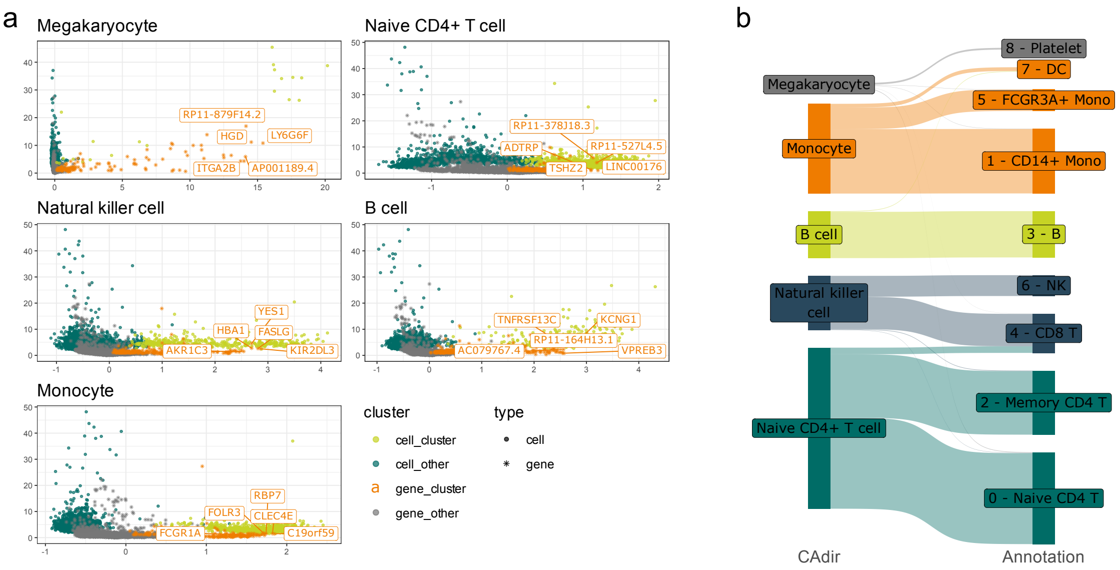


**Suppl. Fig. 2: Clustering of PBMC3k data - additional plots. a**, Association Plots of the corrected clustering of the PBMC3k data after removing outlier cells and genes. Clustered cells are colored in lime green and co-clustered genes in orange. Five genes with the highest $S_\theta$-score are labelled. Other cells and genes are colored in dark green and grey respectively. **b**, Sankey plot comparing the corrected clustering (left) against the annotation obtained through the Seurat vignette (right).

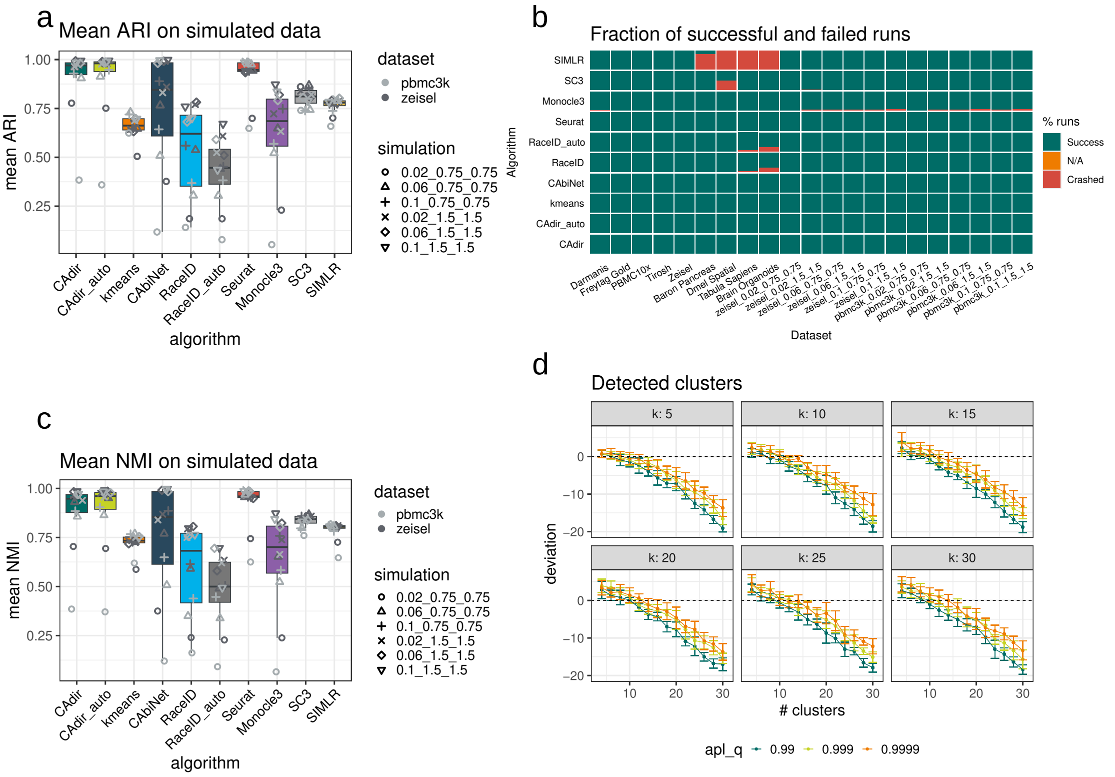


**Suppl. Fig. 3: Average benchmarking results, percentage of successful runs per algorithm and additional cluster detection comparisons.** Mean **a**, Adjusted Rand Index or **c**, Normalized Mutual Information over all 108 parameter combinations for each simulated data set. **b**, Fraction of successful (dark green) runs, runs that produced N/A (not assigned, orange) and crashed runs (red) for each algorithm and tested data set. **d**, Extension of Fig. 4d, with additional quantile cutoffs for the automatic angle determination.
